# Classification of emotions based on functional connectivity patterns of the human brain

**DOI:** 10.1101/2020.01.17.910869

**Authors:** Heini Saarimäki, Enrico Glerean, Dmitry Smirnov, Henri Mynttinen, Iiro P. Jääskeläinen, Mikko Sams, Lauri Nummenmaa

## Abstract

Neurophysiological and psychological models posit that emotions depend on connections across wide-spread corticolimbic circuits. While previous studies using pattern recognition on neuroimaging data have shown differences between various discrete emotions in brain activity patterns, less is known about the differences in functional connectivity. Thus, we employed multivariate pattern analysis on functional magnetic resonance imaging data (i) to develop a pipeline for applying pattern recognition in functional connectivity data, and (ii) to test whether connectivity signatures differ across emotions. Six emotions (anger, fear, disgust, happiness, sadness, and surprise) and a neutral state were induced in 16 participants using one-minute-long emotional narratives with natural prosody while brain activity was measured with functional magnetic resonance imaging (fMRI). We computed emotion-wise connectivity matrices both for whole-brain connections and for 10 previously defined functionally connected brain subnetworks, and trained an across-participant classifier to categorize the emotional states based on whole-brain data and for each subnetwork separately. The whole-brain classifier performed above chance level with all emotions except sadness, suggesting that different emotions are characterized by differences in large-scale connectivity patterns. When focusing on the connectivity within the 10 subnetworks, classification was successful within the default mode system and for all emotions. We conclude that functional connectivity patterns consistently differ across different emotions particularly within the default mode system.

## Introduction

Current neurophysiological and psychological models of emotions largely agree that emotions are characterized by large-scale changes in brain activity spanning both cortical and subcortical areas (see, e.g., Hamann, 2012; Lindquist et al., 2012; Pessoa, 2012; Scarantino, 2012; Adolphs, 2017; Barrett, 2017). Accumulating evidence from both patient and neuroimaging studies in humans show that different brain regions support different aspects of emotional processing, rather than specific emotions *per se* (for reviews, see, e.g., Hamann 2012; Feinstein, 2013; Kragel and LaBar, 2016; Sander et al., 2018; Nummenmaa & Saarimäki, 2019). Accordingly, theoretical frameworks also emphasize the importance of networks of cortical and subcortical regions underlying the representation of emotion in the brain (Pessoa 2017; Barrett & Satpute, 2013; Barrett, 2017).

Multiple lines of evidence suggest that different emotions involve distributed patterns of behavior, and physiological and neural activation patterns. Different categories of emotions have been successfully classified from peripheral physiological responses and bodily activation patterns (Kragel & Labar, 2015; Kreibig et al. 2007; Nummenmaa et al., 2014a; Hietanen et al. 2016; but see Siegel et al., 2018), and subjective feelings (Saarimäki et al., 2016; 2018), suggesting that different emotions have distinguishable characteristics in expressions, bodily sensations and subjective experience (see also review in Nummenmaa & Saarimäki, 2019). At the neural level, multivariate pattern classification studies have revealed that discrete, distributed activity patterns in the cortical midline and somatomotor regions, and subcortical regions such as amygdala and thalamus, underlie different emotion categories (Chikazoe et al., 2014; Wager et al., 2015; Kragel and LaBar, 2015; Peelen et al., 2010; Saarimäki et al., 2016; 2018). Thus, it seems that different instances of the same emotion category share at least some underlying characteristics of neural activity that distinguish the emotions in one category from those in the other. Furthermore, net activity of these regions potentially yields the subjective sensation of the current emotional state, as more similar subjective feelings (such as ‘furious’ and ‘angry’) also trigger more similar neural activation constellations than dissimilar feelings (Saarimäki et al., 2016; 2018).

Yet, as the brain functions as a hierarchy of networks (Bullmore & Sporns, 2009, 2012; Bassett & Sporns, 2017), it is possible that also the connections, rather than local activity patterns, between different regions vary between different emotions (Pessoa 2017; Barrett & Satpute, 2013). In BOLD-fMRI studies, functional connectivity is defined as stimulus-dependent co-activation of different brain regions, usually measured as the correlation of the BOLD time series between two regions. Given the promising results obtained with multivariate pattern analysis of brain activity patterns underlying different emotions, it has been suggested that the study of the brain basis of emotion would benefit from studying the functional relationship between brain regions by employing, for instance, machine learning on functional connectivity patterns (Pessoa, 2018). However, the applications of multivariate pattern analysis in functional networks are sparse. The few studies reporting classification of connectivity measured with fMRI have focused on mental or cognitive states (Richiardi et al., 2011; Shirer et al., 2012; Gonzalez-Castillo et al., 2015) or inter-individual differences in functional connectivity during rest and various tasks (Finn et al., 2015; Shen et al., 2017), demonstrating the plausibility of this approach.

Preliminary evidence highlighting the emotion-related changes in functional connectivity has shown decreased connectivity in specific brain networks after emotional stimulation containing negative affect (Bochardt et al., 2015), and changes in valence and arousal modulate functional connectivity (Nummenmaa et al., 2014). However, the differences in connectivity between different emotion categories have not been previously tested with classification of connectivity patterns calculated from large-scale brain networks. So far, only a handful of studies have compared how specific emotions modulate functional brain connectivity of brain regions, and most of these studies have focused on a limited set of a-priori-defined regions-of-interest (Eryilmaz et al. 2011; Tettamanti et al. 2012; Raz et al. 2016; Huang et al. 2018) or looked at emotion-specific intrinsic connectivity (Touroutoglou et al. 2015). Sustained affective states such as sad mood and anxiety have been found to modulate functional connectivity between the core emotion-related areas such as midline regions including anterior and posterior cingulate cortices, orbitofrontal cortex, insula, and subcortical regions including thalamus and amygdala (Harrison et al., 2008; Seeley et al. 2007; Hermans 2011; 2014; McMenamin et al., 2014). Connectivity especially in salience, frontoparietal control, attention and emotion networks is altered in depression both during generation and regulation of negative emotions as well as during positive emotions (Siegle et al., 2007; Kaiser et al. 2015; Hasler & Northoff 2011; Price et al., 2017). To our knowledge, only one study has investigated whole-brain-level connectivity changes during emotions, but instead of characterizing connectivity patterns across different emotions this study modeled valence- and arousal-dependent connectivity changes (Nummenmaa et al., 2014b). Accordingly, it remains unresolved whether emotion-specific connectivity signatures could distinguish between emotions.

The goals of the current study were two-fold. First, we aimed to demonstrate a pattern classification pipeline as a *proof-of-concept* for studying the functional connectivity patterns underlying different mental states. Second, given the previous studies that have successfully applied machine learning to brain activity patterns underlying different emotions, we tested whether also the functional connectivity patterns during different emotional states could be separated from each other. To reach these goals, we induced six emotions (anger, fear, disgust, happiness, sadness, surprise) and a neutral state using auditory narratives during fMRI scanning. Brain activity related to each emotion was modeled as a functional network. We hypothesized that if distinct connectivity patterns underlie different emotions, the classifier separates between them reliably. The classification was performed with 264 nodes derived from a functional parcellation (Power et al. 2011) and for either whole-brain connectivity (i.e., all nodes together) or for within and between subnetwork connectivity (i.e., for links within and between each subnetwork separately).

## Materials and methods

### Participants

Sixteen female volunteers (ages 20–30, mean age 24.3 years) participated in the fMRI experiment. All participants were right-handed, healthy with normal or corrected-to-normal vision, and gave written informed consent. The studies were run in accordance with the guidelines of the Declaration of Helsinki, and Research Ethics Committee of Aalto University approved the study protocol.

### Stimuli

The stimuli were 35 one-minute-long narratives representing six emotional states (anger, fear, disgust, happiness, sadness, surprise) and a neutral state (five narratives per category). The narratives described personal life events spoken by a female speaker with natural emotional prosody and included emotional expressions, such as weeping and laughing, and have been shown to elicit strong affect in listeners (Smirnov et al., 2019). A separate sample of twenty-four females (ages 20–37, mean age = 24.4 years) rated the stories for the emotional content (Supplementary Figure S1). Behavioral ratings confirmed that the stories successfully elicited the *a priori* defined target emotion; however, to strengthen the effect in the fMRI experiment, the participants also saw a word corresponding the target emotion prior to each story and a short description of the narrative gist (Figure 1a).

**Figure 1.**
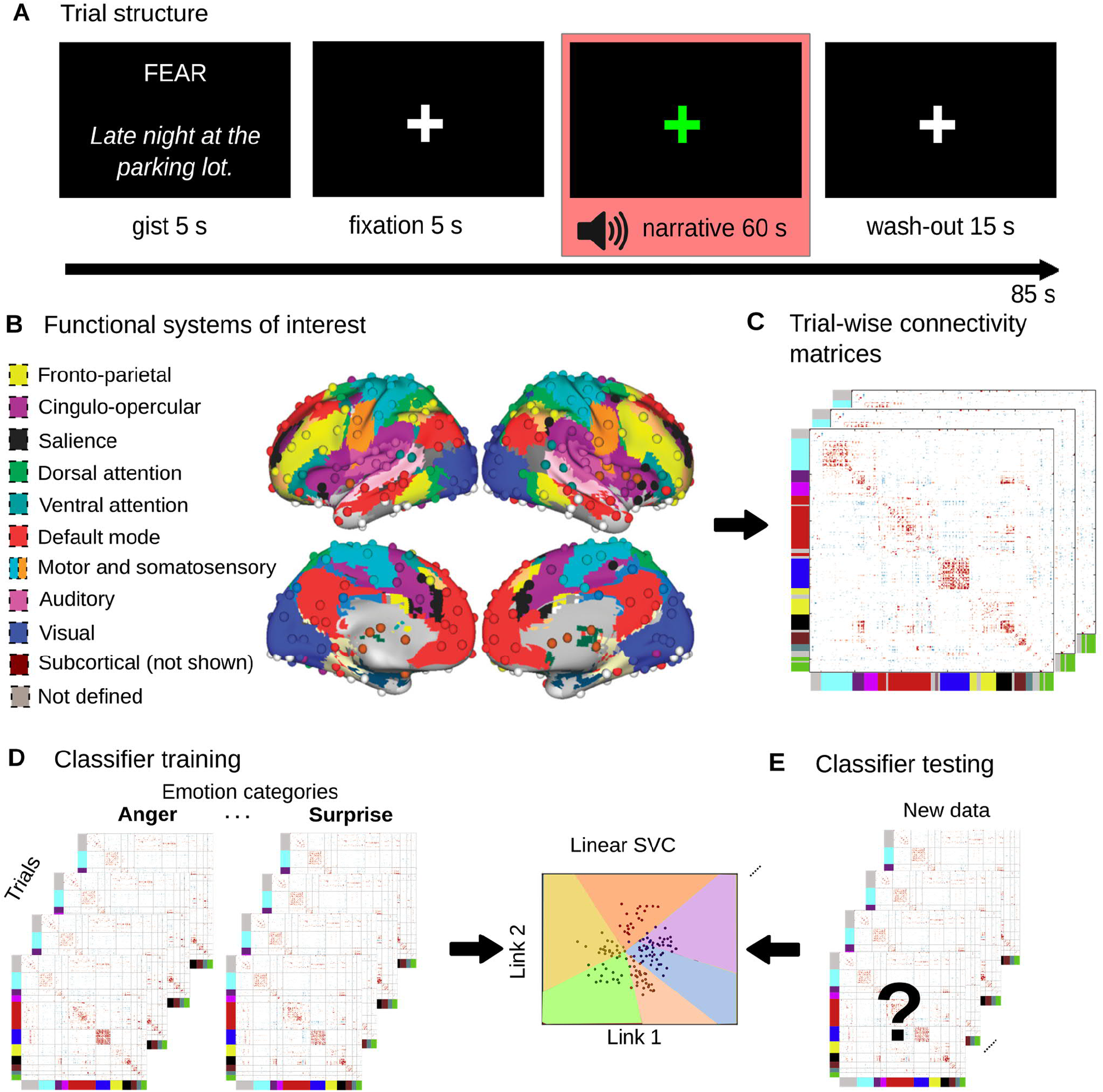
**a)** Trial structure. The highlighted time period (HRF-corrected) was used for calculating the connectivity matrices. **b)** Functional brain systems analyzed in the present study, based on Power et al. (2011). Dots denote network nodes and colors denote subnetworks. **c)** Connectivity matrices were calculated using Pearson correlation between each pair of 264 node time series for each subject and for each 60-s narrative. **d)** The connectivity matrices were fed as input for a linear support vector classifier. **e)** The classifier performance was evaluated by calculating the accuracy (percentage of correct classifier guesses per target category) and the confusion matrix (classifier guesses per category).

### Experimental design

During fMRI, the stimuli were presented in five runs, with one narrative from each emotion category per run. Each run lasted for approximately 10 minutes (365 volumes) and consisted of seven trials (Figure 1a). The order of trials within a run was the same for all participants and the order of runs was counterbalanced across participants (see Supplementary Figure S2). Each narrative was thus presented only once during the experiment. A trial started with a fixation cross presented for 5 seconds, followed by a 5-s presentation of the target emotion (e.g. ‘happy’) and a short description of the narrative gist (e.g. ‘lovers under a tree’). Next, a fixation cross appeared on the screen and the narrative was played through earphones. The trial ended with a 15-s wash-out period allowing emotional state to recover to baseline level following each trial. Subjects were instructed to listen to the narratives similarly as if they would listen to their friend describing a personal life event. The auditory stimuli were delivered through Sensimetrics S14 insert earphones (Sensimetrics Corporation, Malden, Massachusetts, USA). Sound was adjusted for each subject to be loud enough to be heard over the scanner noise. The visual stimuli were back-projected on a semitransparent screen using a 3-micromirror data projector (Christie X3, Christie Digital Systems Ltd., Mönchengladbach, Germany) and from there via a mirror to the participant. Stimulus presentation was controlled with Presentation software (Neurobehavioral Systems Inc., Albany, CA, USA). After the scanning, participants listened to the narratives twice again using an on-line rating tool to continuously rate their subjective valence (ranging from -1 to 1) and arousal (ranging from 0 to 1) during the narrative. Ratings were acquired post-experiment rather than during fMRI, as a reporting task is known to influence neural response to emotional stimulation (Hutcherson et al. 2005; Lieberman et al. 2007) and as repeating a specific emotional stimulus has only a negligible effect on self-reported emotional feelings (Hutcherson et al. 2005).

### Psychophysiological recordings

To remove effects of heart rate and respiration from the BOLD signal during preprocessing, we successfully recorded heart rate and respiration data from 14 subjects with BIOPAC MP150 Data Acquisition System (BIOPAC System, Inc.). Heart rate was measured using BIOPAC TSD200 pulse plethysmogram transducer, which records the blood volume pulse waveform optically. The pulse transducer was placed on the palmar surface of the participant’s left index finger. Respiratory movements were measured using BIOPAC TSD201 respiratory-effort transducer attached to an elastic respiratory belt, which was placed around each participant’s chest to measure changes in thoracic expansion and contraction during breathing. Both signals were sampled simultaneously at 1 kHz using RSP100C and PPG100C amplifiers for respiration and heart rate, respectively, and BIOPAC AcqKnowledge software (version 4.1.1). Respiration and heart rate signals were then used to extract and clean the time-varying heart and respiration rates out of the data with the DRIFTER toolbox (Särkkä et al. 2012).

### MRI data acquisition and preprocessing

MRI data were collected on a 3T Siemens Magnetom Skyra scanner at the Advanced Magnetic Imaging Centre, Aalto NeuroImaging, Aalto University, using a 20-channel Siemens volume coil. Whole-brain functional scans were collected using a whole brain T2*-weighted EPI sequence with the following parameters: 33 axial slices, interleaved order (odd slices first), TR = 1.7 s, TE = 24 ms, flip angle = 70°, voxel size = 3.1 × 3.1 × 4.0 mm^3^, matrix size = 64 × 64 × 33, FOV 198.4 × 198.4 mm^2^. A custom-modified bipolar water excitation radio frequency (RF) pulse was used to avoid signal from fat. High-resolution anatomical images with isotropic 1 × 1 × 1 mm^3^ voxel size were collected using a T1-weighted MP-RAGE sequence.

fMRI data were preprocessed using FSL (FMRIB’s Software Library, www.fmrib.ox.ac.uk/fsl) and inhouse MATLAB (The MathWorks, Inc., Natick, Massachusetts, USA, http://www.mathworks.com) tools (code available at: https://version.aalto.fi/gitlab/BML/bramila). Non-brain matter was removed from functional and anatomical images with FSL BET. After slice timing correction, the functional images were realigned to the middle scan by rigid-body transformations with MCFLIRT to correct subject’s head motion. Next, DRIFTER was used to clean respiratory and heart rate signal from the data (Särkkä et al., 2012). Functional images were registered to the MNI152 standard space template with 2-mm resolution using FSL FLIRT two-step co-registration method with 9 degrees of freedom registration from subject’s space EPI to subject’s space anatomical volume, and 12 degrees of freedom from anatomical to MNI152 standard space. Removal of scanner trend was performed with a 240-seconds long cubic Savitzky-Golay filter (Çukur et al. 2013). To control for head motion artefacts, we followed the procedure as described in Power et al. (2014). The 6 motion parameters were expanded into 24 confound regressors and regressed out. Furthermore, signal at deep white matter, ventricles and cerebrospinal fluid were also regressed out. Finally, temporal bandpass filtering (0.01–0.08 Hz) was applied with the second-order butterworth filter. Spatial smoothing was done with a Gaussian kernel of FWHM 6mm. All subsequent analyses were performed with these preprocessed data.

### Creation of the networks

Because working at voxel-by voxel time series would yield a very high dimensionality and thus be computationally prohibitive, the emotion-specific functional networks were estimated for 264 nodes based on the functional parcellation by Power et al. (2011). We extracted the BOLD time course for each node by averaging the activity of voxels within a 1-cm diameter sphere centered at each node’s coordinates (list of coordinates and module assignments available at http://www.nil.wustl.edu/labs/petersen/Resources_files/Consensus264.xls). For each of the 35 narratives, we calculated the Pearson correlation coefficient between the BOLD time course of each of the nodes during the 60-s-long story, which resulted in a connectivity matrix of 264 × 264 nodes for each narrative (Figure 1b). Next, we removed the baseline connectivity pattern from emotion-wise connectivity matrices by taking the average of the five connectivity matrices for neutral narratives, and regressing it from each of the remaining 30 emotion-specific connectivity matrices separately.

Finally, in addition to the full network of 264 × 264 nodes, we extracted also subnetworks based on the 10 functional systems of interest as proposed by Power et al. (2011; 2014). The subnetworks included were motor and somatosensory (35 nodes), cingulo-opercular (14 nodes), auditory (13 nodes), default mode (58 nodes), visual (31 nodes), fronto-parietal (25 nodes), salience (18 nodes), subcortical (13 nodes), ventral attention (9 nodes), and dorsal attention (11 nodes) networks.

### Correlation between connectivity matrices

To test the similarity between connectivity matrices of different emotions, we quantified the correlation between averaged emotion-wise connectivity matrices with Mantel test employing Spearman correlation as implemented by Glerean et al. (2016; function *bramila_mantel* in https://github.com/eglerean/hfASDmodules/blob/master/ABIDE/bramila_mantel.m). P values were obtained with 5,000 permutations (FDR corrected at p<0.05 for multiple comparisons).

### Classification of connectivity matrices

The classification of emotion categories was performed in Python 2.7.11 (Python Software Foundation, http://www.python.org) using the Scikit learn package (Pedregosa et al. 2011). A between-subjects support vector machine classification algorithm with linear kernel was trained to recognize the correct emotion category out of 6 possible ones (anger, disgust, fear, happiness, sadness, surprise; Figure 1c). Naïve chance level, derived as a ratio of 1 over the number of categories, was 14.2%. The samples for the classifier consisted of the 30 connectivity matrices (5 matrices for each emotion category) from each subject, resulting in altogether 480 samples (80 per category). A leave-one-subject-out cross-validation was performed and the classification accuracy was calculated as an average percentage of correct guesses across all the cross-validation runs (Figure 1d).

For full network classification, the classifier was trained and tested with the full connectivity matrix of each trial. For subnetwork classification, the classifier was trained and tested with the connectivity matrix of each sample either *within* one subnetwork (e.g. connectivity of the nodes within default node network) or *between* two subnetworks (e.g. connectivity between the nodes of default mode network and visual networks, omitting connections of nodes within each network). A separate classifier was trained for each within/between subnetwork division. Based on the subnetwork classifier results (see below), we also investigated the default mode system’s subnetworks in more detail. Therefore, we further split the default mode system into four subnetworks (right temporal, left temporal, midline frontal, and midline posterior) based on clustering of the spatial distances between pairs of nodes within the default mode system and trained a separate classifier for each within/between default subnetworks.

For all classification approaches, we used a permutation test to assess the significance of the results (see, e.g., Combrisson & Jerbi, 2015). To obtain a null distribution, we generated 5,000 surrogate accuracy values for the full network and for each subnetwork separately by shuffling the rows of the upper triangle of the connectivity matrix. The null cumulative distribution function was obtained using kernel smoothing, and the average classification accuracies were compared to the permuted null distribution to obtain their *p* values. Multiple comparisons were corrected for by using the Benjamini-Hochberg FDR correction.

To visualize the differences between connectivity patterns of different emotions, we ran permutation-based t-tests (Glerean 2016) to compare the connectivity matrix of each emotion to that of the rest of the emotions (FDR-corrected for multiple comparisons at p < 0.05). This resulted in a contrast connectivity matrix showing the connections that were unique to each emotion.

## Results

### Classifying emotions from full-brain connectivity patterns

To test whether different emotions are characterized by distinct connectivity patterns, we trained a between-subject classifier with the full network data to recognize the corresponding emotion category out of the six possible ones. Mean classification accuracy was 26% (naïve chance level of 16.6%; permuted *p*<0.00001). The mean classification performance was above the permutation-based significance level for all emotions except sadness (Figure 2A; see Figure 2B for a confusion matrix): anger 23% (FDR corrected p=0.003), disgust 23% (p=0.003), fear 35% (p<0.00001), happiness 28% (p<0.00001), sadness 18% (p=0.291), and surprise 31% (p<0.00001).

**Figure 2.**
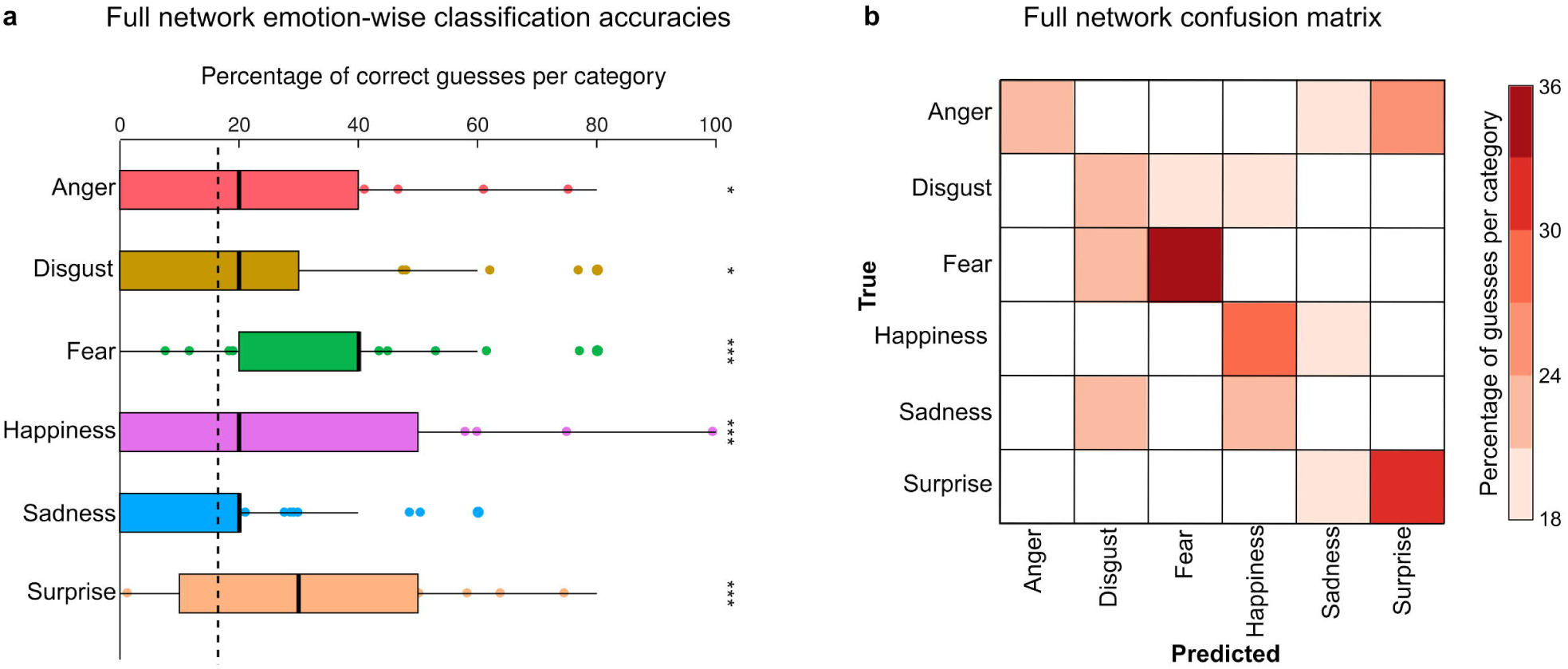
**a)** Emotion-wise classification accuracies for the full-network classification. Dashed line represents naïve chance level (16.6%). Asterisks denote significance relative to chance level (*p<0.01, ***p<0.0001). Thick black line represents median of classification accuracies. Boxes show the 25th to 75th percentiles of classification accuracies and values outside this range are plotted as circles. Whiskers extend from box to the largest value no further than 1.5 * inter-quartile range from the edge of the box. **b)** Classifier confusions from full network classification. Color code denotes average classifier accuracy over the cross-validation runs, cells shown in white have guesses below naïve chance level.

### Classifying emotions from within and between subnetwork connectivity patterns

Our previous studies employing MVPA on brain activity patterns investigated both whole-brain and different regions-of-interest (Saarimäki et al., 2016; 2018). Here we followed a comparable pipeline by implementing the region-of-interest analysis as a *subnetwork-of-interest* analysis. To this end, we separated the connectivity matrices for each functional subnetwork (Power et al., 2014), and trained and tested the across-subject classifiers separately for the connections *within* each subnetwork as well as *between* all possible pairs of subnetworks. Mean classification accuracies and confusion matrices for each within and between subnetwork classifier are shown in Figure 3. Classification accuracy was highest for connections within the default mode system (30%), which after correcting for multiple comparisons, remained the only subnetwork showing significant classification accuracy (p<0.0001; see Supplementary Table S2 for all subnetwork accuracies and p values).

**Figure 3.**
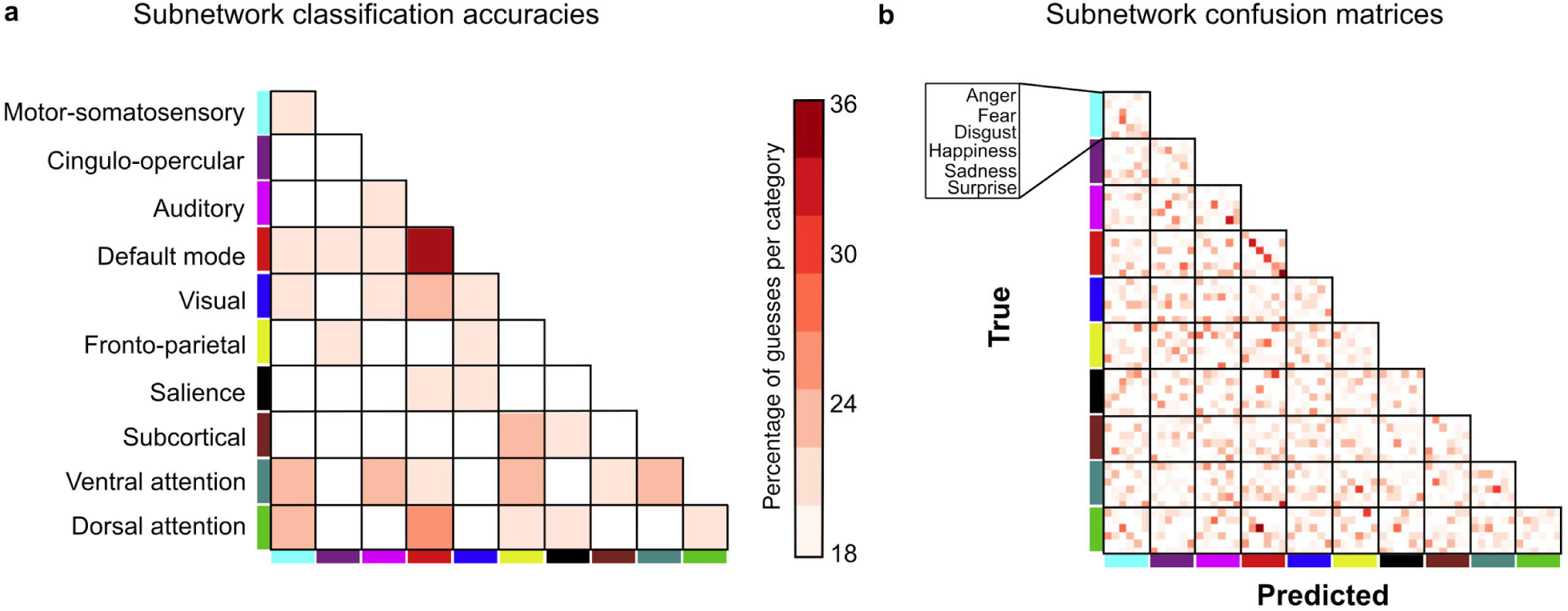
**a)** Classification accuracies for connectivity within and between each ROI. Color code denotes classifier accuracy; cells shown in white have guesses below naïve chance level (16.6%). After correcting for multiple comparisons, only the accuracy for within default mode network connections remained significant. **b)** Classifier confusions for subnetwork classification.

To visualize the emotion-specific functional connectivity, we plotted the connectivity matrices (Supplementary Figure S4; see Supplementary Figure S3 for connectivity matrix of the neutral state). Correlations between each pair of connectivity matrices for all emotions and the neutral state were significant (Supplementary Figure S4; correlations ranging from rho=0.79–0.84, all Bonferroni-corrected Ps<0.01; see Supplementary Table S1). The average connectivity patterns for each emotion with neutral baseline removed are shown in Supplementary Figure S5 and pairwise t-tests between the connectivity matrices in Supplementary Figure S7.

### Classifying emotions from default mode network connectivity

Because default mode system was the only subnetwork where connectivity-based classification accuracies were above chance-level after post-hoc correction, we next investigated its connectivity patterns in more detail. Within the default mode system, all emotions could be classified with above-chance-level accuracy (see Figure 4a): anger 24% (p=0.001), disgust 33% (p<0.00001), fear 30% (p<0.00001), happiness 31% (p<0.00001), sadness 26% (p=0.0002), and surprise 35% (p<0.00001). See Supplementary Figure S6 for emotion-specific connectivity patterns for default mode network.

**Figure 4.**
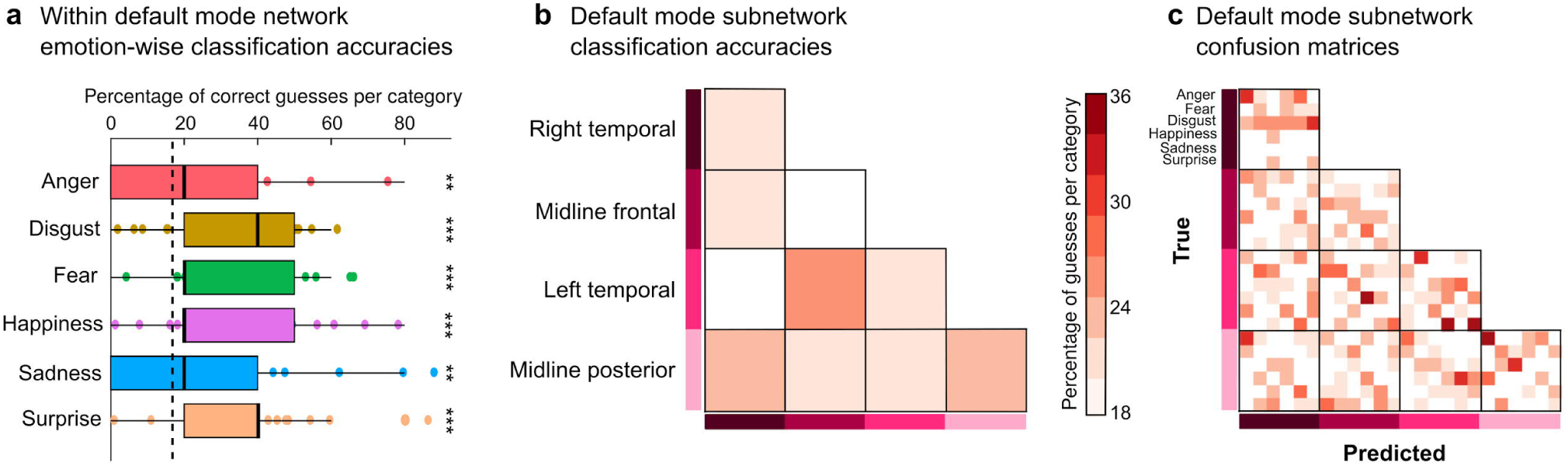
**a)** Emotion-wise classification accuracies for connections within the default mode system. Dashed line represents naïve chance level (16.6%). Asterisks denote significance relative to chance level (**p<0.001, ***p<0.0001). Thick line represents median of classification accuracies. Boxes show the 25th to 75th percentiles of classification accuracies and values outside this range are plotted as dots. Whiskers extend from box to the largest value no further than 1.5 * inter-quartile range from the edge of the box. **b)** Classification accuracies and **c)** subnetwork confusion matrices for DMN subnetwork classification. Color code denotes classifier accuracy; cells shown in white have guesses below naïve chance level.

We next investigated whether there are node-wise differences in classification accuracy within the default mode network by splitting DMN nodes to four subnetworks based on their spatial proximity in the brain. The spatial clustering of DMN resulted in four DMN subnetworks: left temporal, right temporal, frontal, and posterior midline. We trained classifiers to recognize emotions from the connections within and between these DMN subnetworks separately. Classification was successful for connections within midline posterior subnetwork (22.5%, permuted and FDR-corrected p=0.008), between left temporal and midline frontal subnetworks (23.5%, p<0.001), and between right temporal and midline posterior subnetworks (21.5%, p=0.032). The classification accuracies and confusion matrices are shown in Figure 4.

## Discussion

The goals of the current study were to (i) provide a *proof-of-concept* for the application of machine learning to functional connectivity patterns, and to (ii) test whether different emotional states can be classified from functional connectivity. We show differences in whole-brain functional connectivity patterns during different emotional states (anger, fear, disgust, happiness, and surprise), as evidenced by significantly above chance-level classification accuracy. The classifier was trained across participants, demonstrating that the connectivity patterns were consistent across subjects. Emotion-specific connectivity patterns were most prominently observed within the default mode system.

The present results thus show that not only regional activity patterns (Saarimäki et al., 2016; Kragel & LaBar 2016), but also large-scale connectivity changes across specific brain systems, underlie different emotional states. The overall accuracy (26%) in the current study is above chance (here, 16.6% for the six emotion categories), yet clearly smaller than the 50–60% accuracies reached by regional classifiers trained to classify an equal number of emotional states (Saarimäki et al., 2016). Furthermore, while previous studies have shown successful classification of connectivity underlying different mental or cognitive states, such as resting and movie viewing (Richiardi et al., 2011; Shirer et al., 2012; Gonzalez-Castillo et al., 2015), the current study shows that we can to some extent classify varying emotional states evoked by same cognitive task (mental imagery of narratives) using large-scale brain networks. There are multiple possible reasons for this discrepancy in the classifier performance but it most likely reflects the low general power on fMRI connectivity analysis due to slow-frequency signal of interest. Future studies should address this issue using, for example, MEG experiments yielding better temporal resolution for dynamic connectivity measures. Moreover, such classification of emotions focusing on a smaller set of connections between specific regions of interest has not been successful (Raz et al., 2016). Therefore, the current work proposes that especially large-scale connectivity is modulated by different emotional states to the extent that classification of connectivity patterns between individuals is possible. While multivariate pattern analysis results are not straightforward to interpret in a neurophysiological sense, they might reflect some kind of net activation of the system that is picked up by the classifier (see also Nummenmaa & Saarimäki, 2019).

### Functional connectivity is modulated differently by different emotions

Our main new finding was that different whole-brain connectivity patterns underlie specific emotional states, similarly as has previously been found for regional activity patterns (for reviews, see Kragel & LaBar 2016; Nummenmaa & Saarimäki, 2019). We have previously shown that anger, fear, disgust, happiness, sadness, and surprise can be classified from brain activity in the cortical midline regions, subcortical regions, somatomotor regions, but also regions related to cognitive functions such as memory and language (Saarimäki et al. 2016), thus suggesting that different emotions modulate each of these regions differently. These areas are also consistently activated in studies using univariate analysis of emotional brain responses (Kober et al. 2008). Altogether these results thus suggest that an individual’s emotional state is based on the net activation of all these regions (see also Nummenmaa & Saarimäki, 2019), and that the resultant conscious experience or subjective emotional feeling may arise from activity in the cortical midline regions where information from different parts of the brain is integrated (Northoff et al., 2006, Damasio et al., 2000; Saarimäki et al. 2016). Together, the results from the voxel-based and connectivity-based pattern classification of emotions support a model where emotions are brought about by distributed net activation of different regions together with their connectivity patterns.

As highlighted by the Figures 2A and 4B, classification accuracy varied across cross-validation runs. With our leave-one-participant-out classifier, this means that there is variation in how well the connectivity patterns of different individuals can be classified. Addressing the individual differences in this classification performance is out of the scope of the current paper but constitutes an interesting topic for future research. To further investigate which connections might differ between emotions, we trained separate classifiers for connections within and between different functional subnetwork. Interestingly, classification accuracy was significantly above chance level only within the default mode system connectivity suggesting that connectivity differences between emotions stem mainly from these connections. This warrants a closer look at where these differences within DMN stem from.

### Emotion-specific differences in default mode network connectivity

Connectivity-based classification accuracy was highest when we considered only the connections within the default mode system. This is in line with the amounting evidence that shows the importance of default mode system regions in emotional processing. These areas contain the most robust distinct voxel activity patterns for different emotions (Saarimäki et al. 2016; Wager et al., 2015; Kragel & LaBar 2015) and have been suggested to serve in integrating information about one’s internal state (Klasen et al., 2011; Mar 2011; Kleckner et al., 2017) and hold representations for self-relevant mentalization (Summerfield et al., 2009; D’Argembeau et al. 2010; Andrews-Hanna et al. 2014). The activity patterns resulting from the integration of different representations, such as those of salience, potential motor actions, bodily sensations, or sensory features, may constitute a core feature of the emotional state, with distinct signatures for different emotions (Saarimäki et al., 2016). In previous studies, changes in DMN connectivity during emotional processing have been found during sad mood (Harrison et al. 2008) and depression (Whitfield-Gabrieli & Ford, 2012), and emotional stimulation continues to affect the DMN connectivity minutes after the stimulus ends (Eryilmaz et al., 2011), suggesting that these regions and their connectivity play a role in *sustaining* an emotional state.

Default mode network can be further divided into subsystems that serve different cognitive functions (Buckner et al., 2008; Andrews-Hanna et al. 2010, 2014). Therefore, we examined whether emotions can be classified from connectivity patterns within and between different DMN subsystems. In our spatial parcellation, the default mode subnetworks comprised medial PFC, posterior midline regions, and left and right lateral temporal areas (Power et al. 2011). We found that emotions could be classified especially from connectivity patterns within posterior midline DMN connections. Midline posterior regions including posterior cingulate cortex and precuneus have been linked to integration of information from other DMN subsystems and other brain regions (Buckner et al., 2008; Andrews-Hanna et al. 2014). Successful classification of emotions from the connectivity patterns in this region suggest that emotions might vary in how and which information is integrated during the emotional state. For instance, interpretation of emotions with a strong action component (e.g., fear) might rely more on motor inputs, while the role of such inputs is smaller with emotional states that do not require immediate motor actions (e.g., sadness). This accords with the view that emotions arise from integrated activity across multiple physiological, behavioural and neural systems (Nummenmaa & Saarimäki, 2019).

Furthermore, emotions could be classified also from connections between midline frontal and left lateral temporal regions, and between midline posterior and right lateral temporal regions. Frontal midline regions including medial PFC are related to the construction of self-relevant mental simulations (Buckner et al., 2008), whereas lateral temporal lobes are related to conceptual processing and storing of semantic and conceptual information (Patterson et al., 2007; Binder & Desai, 2011). Differences between the connectivity of these regions suggest that emotions differ in how conceptual information is integrated in the construction of self-relevant mentalization in mPFC and integrated with other types of information in the PCC. However, it is noteworthy that none of the subnetwork connections exceeded the classification accuracy of all connections within the default mode system, suggesting that emotion-wise differences are strongest when looking at the DMN functional connectivity as whole.

We found no significant classification performance for the connectivity within or between areas other than default mode network despite the evidence showing that activation patterns within a large number of other areas, including subcortical, somatomotor, and frontal areas (Saarimäki et al., 2016), differ between emotions. This is probably partly due to how connectivity was calculated in the current study: connectivity was calculated over a time period of one minute, which is less sensitive to rapid temporal changes in connectivity that potentially underlie different emotional states (see, e.g., Pessoa, 2018). For instance, Nummenmaa et al. (2014b) have shown that such dynamic changes reveal large connectivity differences in positive versus negative valence and high versus low arousal. Further, it is unlikely that emotional state remains the same over the 1-minute-long stimulation, probably restricting the stability of connectivity patterns and affecting their distinctness. However, connectivity measures with shorter time windows usually contain more noise which is why, for instance, resting state connectivity is measured for preferably over 5 minutes. Thus, classification of connectivity patterns in general is a more difficult task than that of activation patterns; yet, the current proof-of-concept work shows that it is possible.

The current experiment had a 15 second wash-out period between emotional stimuli. It is likely that emotion effects from the previous trial are still present in the following trial, as it has been shown that emotional induction lasts for minutes after the stimulation ends (Eryilmaz et al., 2010). However, the experimental stimulation of the next trial begins immediately after the wash-out period, allowing new emotional stimulus to modulate the emotion systems and likely overriding the previous input. Moreover, the problem was alleviated by varying the order of emotions between trials.

### Conclusions

We conclude that basic emotions including anger, fear, disgust, happiness, sadness, and surprise differ in the connectivity patterns across the brain in a manner that is consistent across individuals. Connectivity-based classification of emotions was most accurate within the default mode system, suggesting that connectivity across this region contains the most accurate representation of the individual’s current emotional state. Together with previous regional voxel-based pattern classification results, the present findings support the view that emotions are represented in a distributed fashion across the brain and the current emotional state is defined by the net activity of the system.

## Acknowledgements

This work was supported by Academy of Finland (#265917 to L.N., and #138145 to I.P.J.), ERC Starting Grant (#313000 to L.N.); Finnish Cultural Foundation (#00140220 to H.S.); Kordelin Foundation (#160387 to H.S.); and Russian Federation Government (#075-15-2019-1930 to I.P.J). We thank Marita Kattelus for her help with the data acquisition. We acknowledge the computational resources provided by the Aalto Science-IT project.

