## Supplementary Material for "Classification of emotions based on functional connectivity patterns of the human brain"

### Supplementary Figure Legends

**Figure S1.** The stimuli consisted of 35 one-minute-long narratives that induced six emotional states and a neutral state. A sample of 24 female participants not included in the fMRI experiment rated on a scale from 0 to 5 how much of each emotion was elicited by the narrative. The coloring indexes the mean intensity for experiencing each emotion for each narrative.

**Figure S2.** The order of the narratives. The order of the trials within a run was the same for all participants. The order of the runs was counterbalanced across participants.

**Figure S3.** Average connectivity matrix for the neutral state.

**Figure S4.** Mean connectivity matrix for each emotion. Line strength between matrices shows the Spearman correlation between each matrix pair. Strength of correlations is denoted by line color and width. Data are thresholded at $p < 0.05$, FDR corrected for multiple comparisons.

**Figure S5.** Connectivity matrices for each emotion with neutral baseline removed.

**Figure S6.** Connectivity matrices of within default mode network connections for each emotion with neutral baseline removed.

**Figure S7.** Pairwise t tests showing the differences in full network connectivity for each emotion contrasted with all other emotions. P values corrected for multiple comparisons using FDR correction ($p<0.05$).

### Supplementary Table Legends

**Table S1.** Correlation between each pair of average connectivity matrices. Correlation was calculated using Spearman correlation coefficient. P values were obtained with permutations and FDR corrected to account for multiple comparisons. All correlations were significant at $p<0.00001$.

**Table S2.** Classification accuracies and permuted p values (FDR corrected for multiple comparisons) for subnetwork classification.

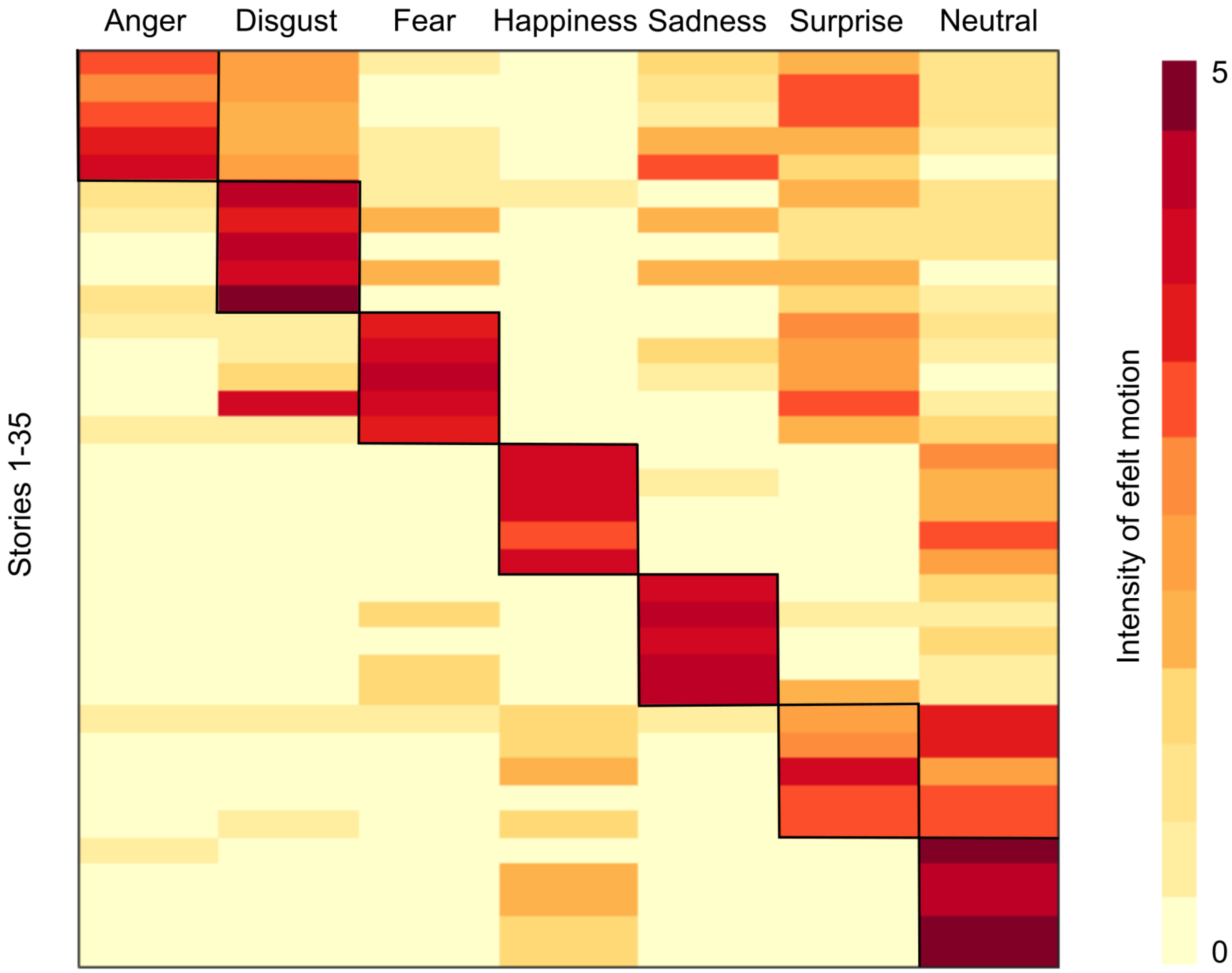


**Figure S1.** The stimuli consisted of 35 one-minute-long narratives that induced six emotional states and a neutral state. A sample of 24 female participants not included in the fMRI experiment rated on a scale from 0 to 5 how much of each emotion was elicited by the narrative. The coloring indexes the mean intensity for experiencing each emotion for each narrative.

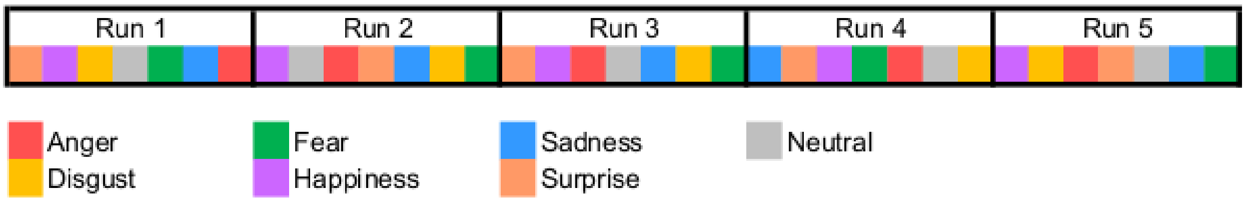


**Figure S2.** The order of the narratives. The order of the trials within a run was the same for all participants. The order of the runs was counterbalanced across participants.

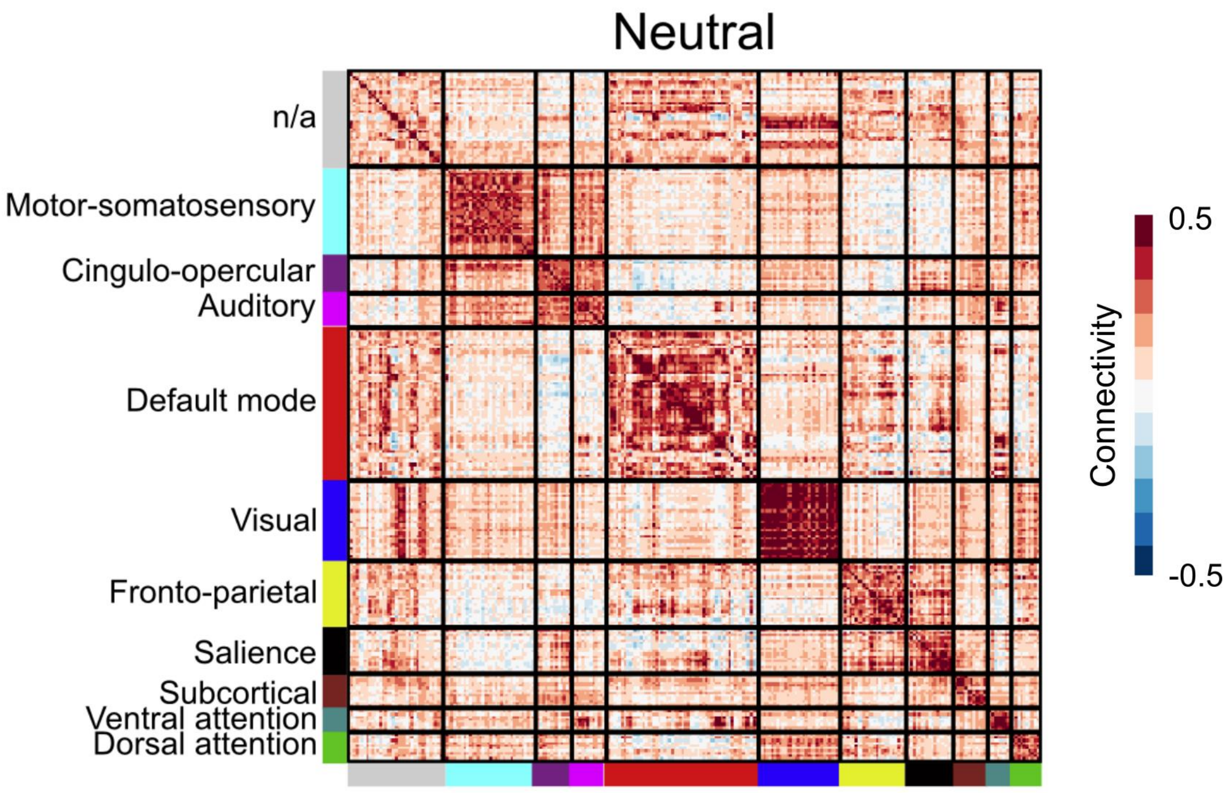


**Figure S3.** Average connectivity matrix for the neutral state.

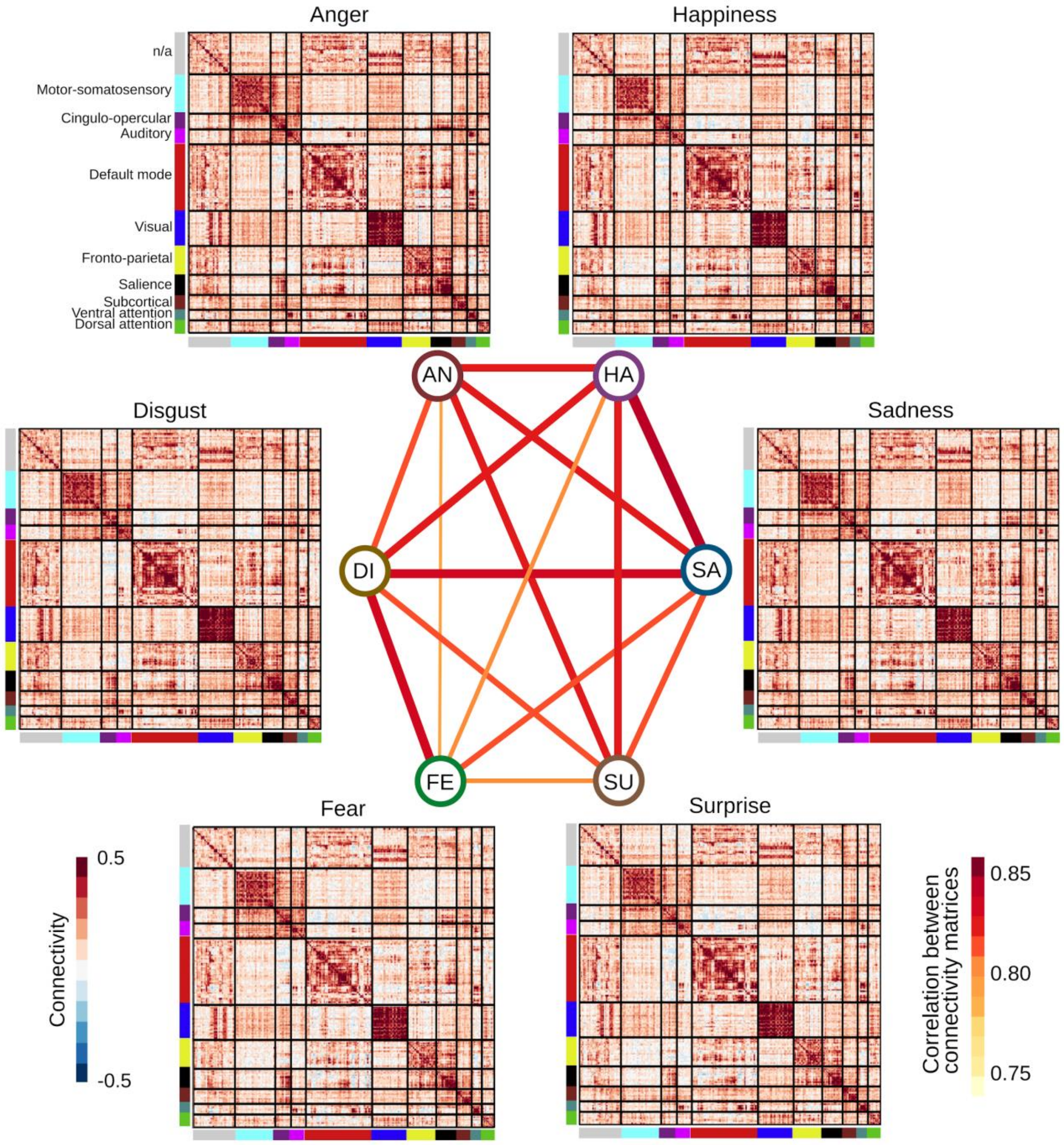


**Figure S4.** Mean connectivity matrix for each emotion. Line strength between matrices shows the Spearman correlation between each matrix pair. Strength of correlations is denoted by line color and width. Data are thresholded at $p < 0.05$, FDR corrected for multiple comparisons.

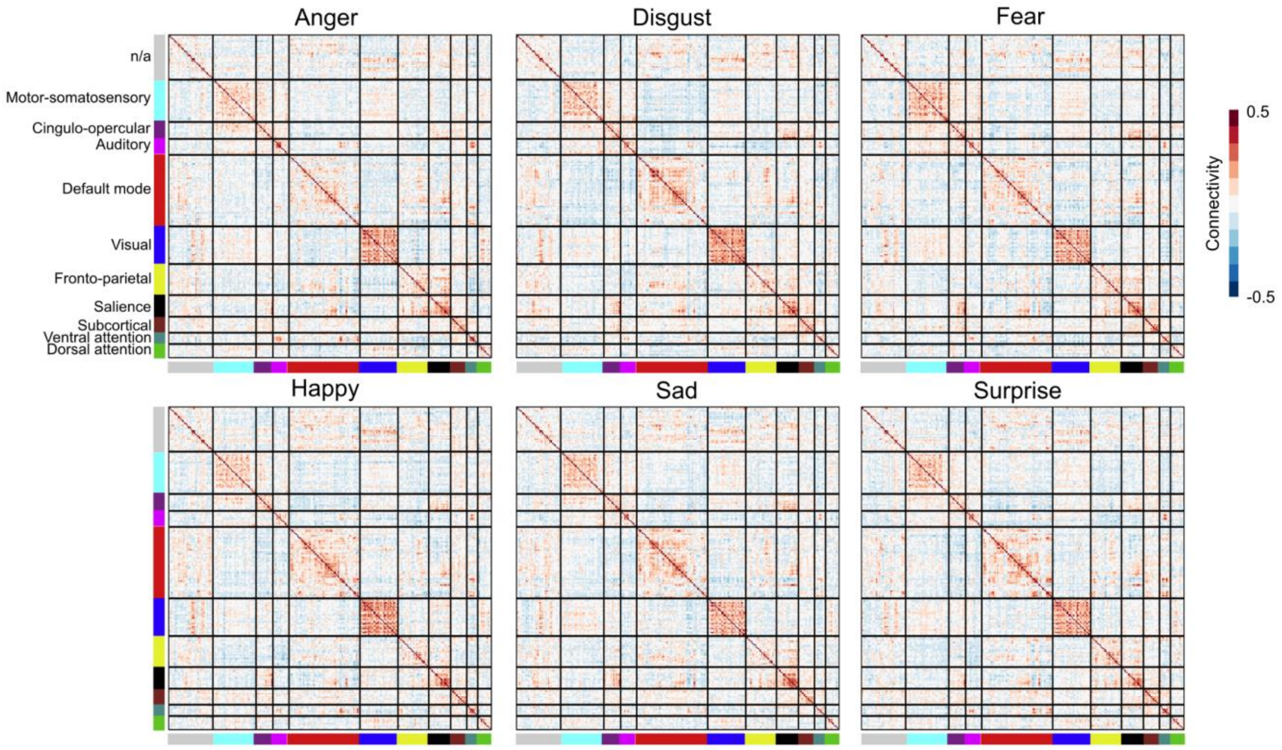


**Figure S5.** Connectivity matrices for each emotion with neutral baseline removed.

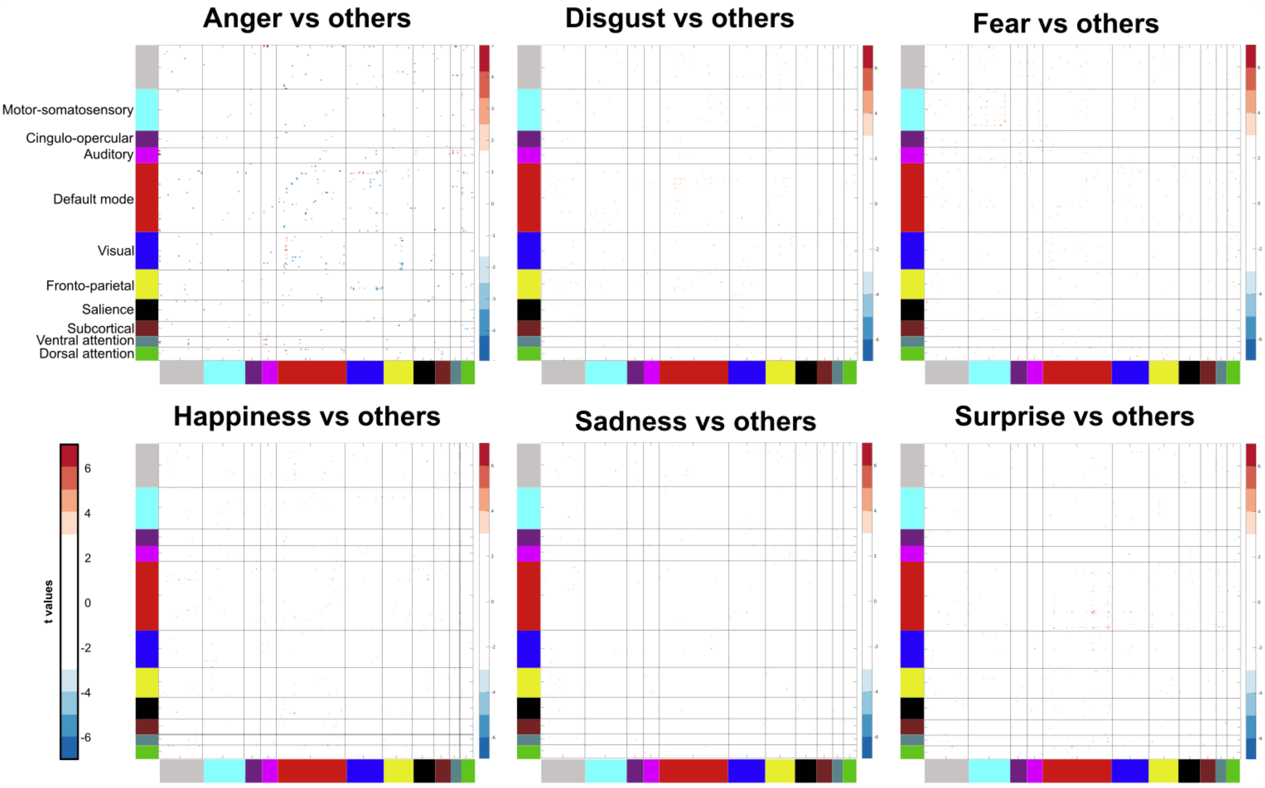


**Figure S6.** Connectivity matrices of within default mode network connections for each emotion with neutral baseline removed.

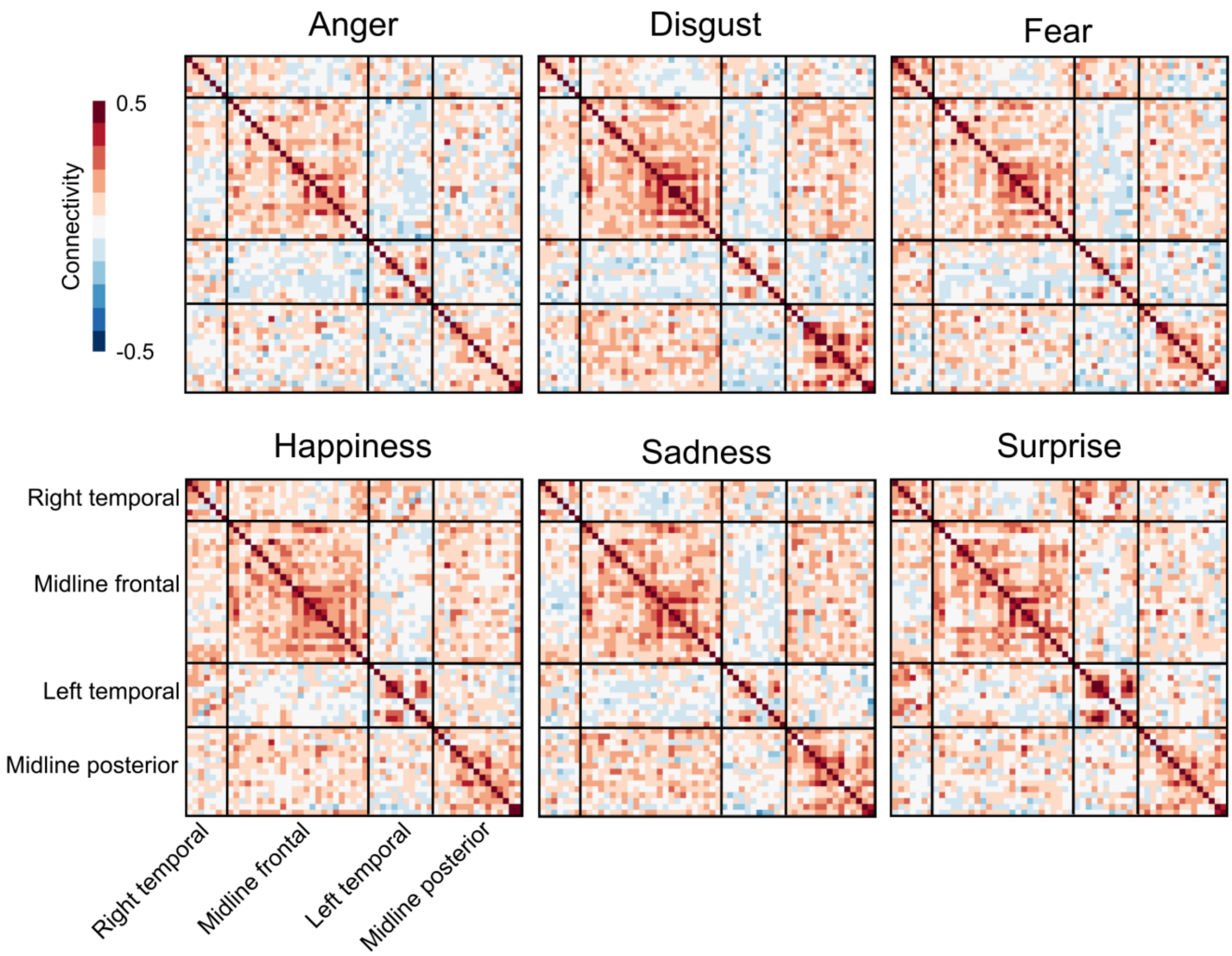


**Figure S7.** Pairwise t tests showing the differences in full network connectivity for each emotion contrasted with all other emotions. P values corrected for multiple comparisons using FDR correction ($p<0.05$).

**Table S1.** Correlation between each pair of average connectivity matrices. Correlation was calculated using Spearman correlation coefficient. P values were obtained with permutations and FDR corrected to account for multiple comparisons. All correlations were significant at $p<0.00001$.

| | Disgust | Fear | Happy | Sad | Surprise | Neutral |
|---|---|---|---|---|---|---|
| Anger | 0,81 | 0,79 | 0,82 | 0,82 | 0,82 | 0,83 |
| Disgust | | 0,83 | 0,82 | 0,83 | 0,81 | 0,81 |
| Fear | | | 0,80 | 0,81 | 0,80 | 0,79 |
| Happy | | | | 0,84 | 0,82 | 0,84 |
| Sad | | | | | 0,81 | 0,81 |
| Surprise | | | | | | 0,81 |

**Table S2.** Classification accuracies and permuted p values (FDR corrected for multiple comparisons) for subnetwork classification.
